# Dual phage sensing activates an RNA-targeting reverse transcriptase for anti-phage defense

**DOI:** 10.64898/2026.10.27.754916

**Authors:** Xin-Yang Wei, Hua Qi, Xin-Yi Song, Linshu Li, Yushan Xia, Jun-Tao Zhang, Ning Jia

## Abstract

Defense-associated reverse transcriptases (DRTs) mediate diverse antiphage responses, yet how they sense phage infection and couple unconventional DNA synthesis to antiviral effector activation remains poorly understood. Here, we characterize DRT4 as a dual-sensor immune system that integrates two distinct phage-derived signals. Cryo-EM analysis reveals that DRT4 assembles into a dimer-of-trimers hexamer, with each protomer catalyzing protein-primed, template-independent synthesis of a covalently linked ssDNA product. DRT4 displays a strong preference for purine nucleotides, determined by base-specific interactions within the polymerase active site. Using phage escape genetics, we identify two independent activators of DRT4: the phage ribonucleotide reductase NrdAB, which increase intracellular dNTP levels, and the SSB-like protein 0087, which directly binds the DRT4-generated ssDNA. Activated DRT4 exhibits dNTP-dependent RNase activity, linking phage sensing to RNA cleavage and abortive infection. Together, our findings show DRT4-generated ssDNA as a signaling intermediate that integrates phage-induced metabolic changes with direct recognition of viral replication proteins, revealing how protein-primed DNA synthesis can be repurposed for infection sensing and antiviral effector activation.

## Introduction

Reverse transcriptases (RTs), found across all three domains of life, are defined by their ability to synthesize DNA from RNA templates^1^. Beyond their canonical roles in retroelements and genome evolution, RTs have recently emerged as remarkably diverse components of bacterial antiviral immunity^2,3^. Antiviral RTs occur in CRISPR-associated systems, retrons, abortive infection (Abi) systems, and a broad group of defense-associated reverse transcriptases (DRTs) encoded within genomic defense islands^3–5^. DRTs encompass several phylogenetically and mechanistically distinct families and can be broadly classified into three major classes^2^. Class 1 DRTs are typically fused to a C-terminal α-helical repeat (αRep) domain and frequently catalyze template-independent DNA synthesis. Class 2 DRTs are commonly associated with structured non-coding RNAs (ncRNAs) that serve as templates for DNA synthesis, whereas Class 3 DRTs are often genetically linked to predicted enzymatic effectors, suggesting functional coupling between reverse transcription and downstream effector activation^2^. Despite their widespread distribution, how DRTs detect phage infection and convert unconventional DNA synthesis into antiviral activity remains incompletely understood.

Recent studies of Class 2 DRTs have revealed strikingly diverse strategies for coupling reverse transcription to immunity. In the DRT2 system, reverse transcription of an ncRNA proceeds through iterative template jumping to generate long concatemeric cDNA (ccDNA)^6–8^. This cDNA contains a de novo open reading frame, *neo*, whose translation produces a toxic Neo protein directly targets the bacterial membrane, inducing membrane depolarization and growth arrest to restrict phage propagation^6–8^. Our recent work further showed that DRT2 can sense two features associated with active phage replication: changes in nucleotide availability driven by the phage ribonucleotide reductase NrdAB and direct recognition of phage single-stranded DNA-binding or single-strand annealing proteins (SSB/SSAP) through the initially synthesized DNA product^8^. Thus, the nascent DNA generated by DRT2 acts not only as an intermediate of reverse transcription but also as a platform for infection sensing. DRT9 employs a distinct strategy. Rather than producing a protein-coding cDNA, DRT9 synthesizes long poly(A)-rich single-stranded DNA in response to phage-induced changes in nucleotide metabolism^9,10^. The resulting DNA acts as a molecular decoy that captures phage SSB proteins and thereby disrupts viral DNA replication^9^. DRT3 combines ncRNA-templated poly(TG) synthesis by DRT3A (a Class 2-type UG3 RT) with protein-primed, template-independent poly(CA) synthesis by DRT3B (a Class 1-type UG8 RT) to generate a self-complementary DNA product, although its activation and effector mechanisms remain incompletely understood^11,12^. DRT10 comprises an RT, ncRNA, and SLATT effector, and generates 7-nt tandem-repeat ssDNA through protein-primed ncRNA-templated synthesis; phage SSB-like proteins promote activation, whereas Host SbcB limits cDNA accumulation^13,14^.

Moreover, Class 3 system DRT1 are proposed to use protein-primed, template-free synthesis of semirandom ssDNA to regulate its nitrilase effector between a dormant filamentous state and an activated state^15^. DRT11, another Class 3 system, also requires nitrilase activity for its defensive function^16^. Together, these findings illustrate that DRT-generated DNA can function as a coding intermediate, molecular decoy, infection sensor, or allosteric regulator rather than simply as a conventional reverse-transcription product.

Class 1 DRTs remain mechanistically less understood. AbiA, AbiK and Abi-P2 comprise a distinct group of RT-like antiviral proteins that use protein-primed, template-independent DNA synthesis rather than canonical reverse transcription^17–20^. Structural studies reveal a conserved RT-like domain coupled to a characteristic helical domain, while phage escape from AbiK frequently occurs through mutations in viral single-strand annealing proteins^21^, suggesting a functional connection between RT-generated ssDNA and phage replication machinery. DRT4 is a representative Class 1 system containing an N-terminal RT-like domain and a C-terminal α-helical domain^2^. We therefore set out to investigate how DRT4 senses physiological phage signals and converts this recognition into antiviral defense.

Here, we show that DRT4 forms a dimer-of-trimers hexamer in which each protomer catalyzes protein-primed, template-independent synthesis of purine-rich ssDNA. Through phage escape genetics, we identify two distinct phage-derived activators of DRT4: the ribonucleotide reductase NrdAB, which alters intracellular dNTP availability, and the SSB-like protein 0087, which directly binds the DRT4-generated ssDNA, linking recognition of a phage replication protein to activation of the downstream effector state. Together, our findings establish DRT4-generated ssDNA as a signaling intermediate that integrates phage-induced metabolic changes with direct signals of active phage replication to trigger RNA cleavage and abortive infection.

## Results

### DRT4 confers anti-phage defense via abortive infection

To investigate the mechanism of DRT4-mediated anti-phage defense, we cloned the DRT4 gene from *Escherichia coli* 21-C8-A into *E. coli* MG1655 and challenged the resulting strain with a diverse panel of phages from the BASEL collection (Figure 1A). DRT4 conferred robust protection against a broad spectrum of phages, with particularly strong activity against members of the *Dhillonvirus*, *Demerecviridae*, *Queuovirinae*, and *Vequintavirinae* groups, reducing efficiency of plating (EOP) by up to ∼10^6^-fold (Figure 1B). Protection against representative phages was pronounced at low multiplicities of infection but was largely overcome at a high MOI of 5 (Figure 1C and Figure S1), a phenotype characteristic of abortive infection (Abi) systems^22^. DRT4 did not affect phage adsorption to host cells (Figure 1D), but instead acted at a post-adsorption stage to suppress phage replication and progeny production (Figure 1E). Consistently, time-resolved colony-forming unit (CFU) measurements showed that DRT4-expressing cells failed to recover following high-MOI infection, indicating that activation of DRT4 triggers host cell death to limit phage propagation (Figure 1F).

**Figure 1.**
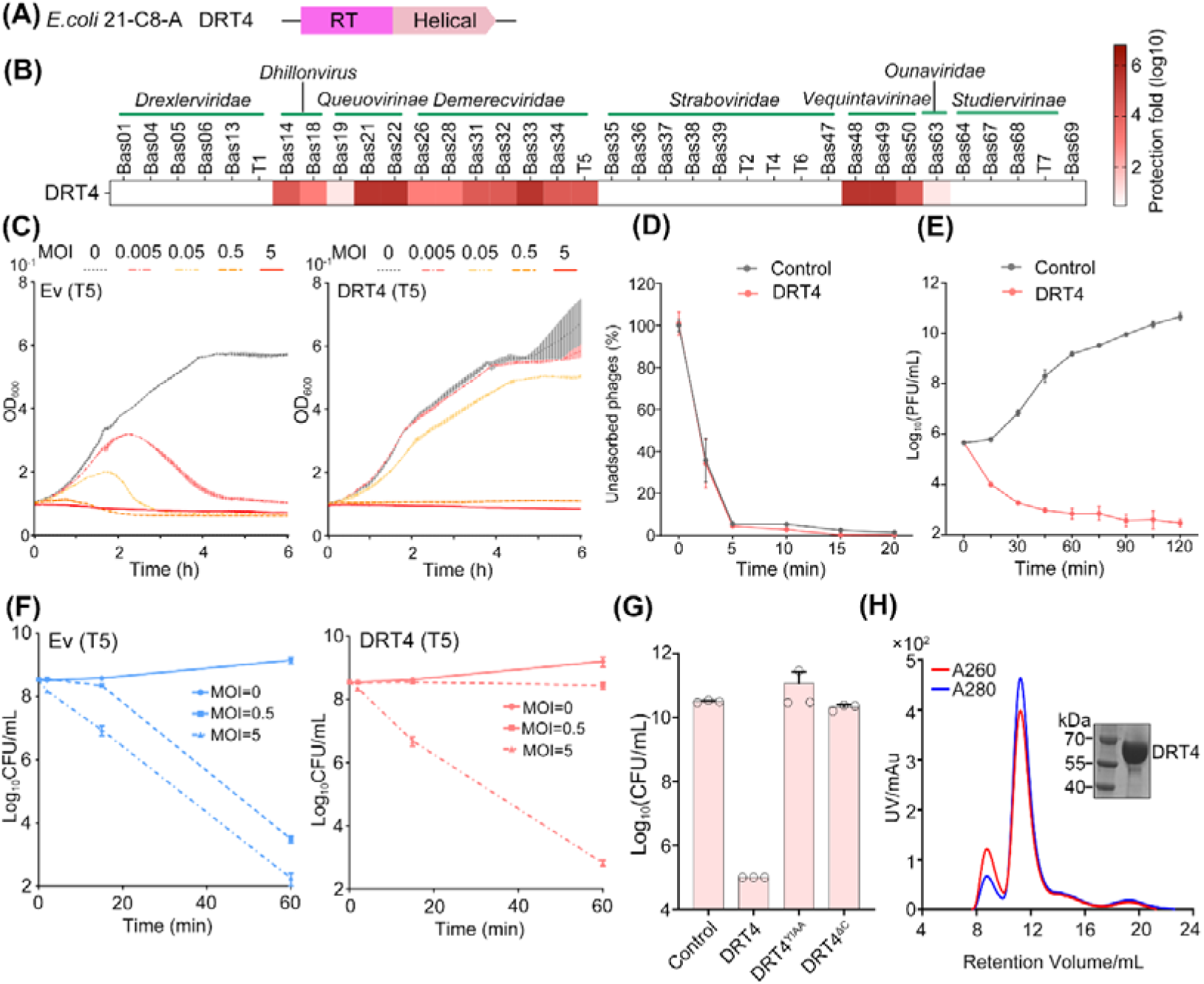
| DRT4 defend against phage infection through abortive infection. **(A)** Schematic of the DRT4 locus from *E. coli* strain 21-C8-A. **(B)** Plaque assays showing phage susceptibility of *E. coli* MG1655 carrying DRT4 or an empty vector (EV) against the indicated phages. Images are representative of three biological replicates. **(C)** Growth curves of *E. coli* MG1655 carrying DRT4 or EV following infection with phage T5 at the indicated multiplicities of infection (MOIs). Phage was added at time 0. **(D)** Adsorption of phage T5 to *E. coli* MG1655 carrying DRT4 or EV. **(E)** One-step growth curves of phage T5 in *E. coli* MG1655 carrying DRT4 or EV. **(F)** Survival of *E. coli* MG1655 carrying DRT4 or EV following 60 min of T5 infection at the indicated MOIs, measured by colony-forming unit (CFU) counts. **(G)** Anti-phage activity of wild-type DRT4 and the indicated mutants. *E. coli* MG1655 carrying wild-type DRT4, DRT4^YIAA^, DRT4^Δhelical^, or EV was challenged with phage T5. Data are shown as the mean of three biological replicates, with individual data points overlaid. **(H)** Size-exclusion chromatography profile of the purified DRT4–ssDNA complex. Inset, SDS–PAGE analysis of the indicated fractions.

DRT4 comprises an N-terminal reverse transcriptase (RT) domain followed by a C-terminal helical domain (Figure 1A). To determine the contribution of these domains to antiviral defense, we substituted the two conserved aspartate residues within the YIDD catalytic motif (Y238–D241) of the RT domain with alanine and separately deleted the C-terminal helical domain. Both perturbations abolished anti-phage activity (Figure 1G), demonstrating that RT catalytic activity and the C-terminal domain are both essential for DRT4-mediated defense. Size-exclusion chromatography (SEC) of purified DRT4 further revealed a prominent high-molecular-weight species with an A260/A280 ratio of ∼0.8, higher than expected for protein alone, suggesting that DRT4 forms a stable oligomeric complex associated with tightly bound nucleic acids that persist throughout purification (Figure 1H). Together, these findings indicate that DRT4 forms an oligomeric nucleoprotein defense complex that restricts phage propagation through abortive infection by triggering host cell death, with its RT catalytic activity being indispensable for anti-phage defense.

### DRT4 adopts a dimer-of-trimers hexameric architecture with each protomer bound a protein-primed ssDNA

To elucidate the molecular basis of DRT4-mediated antiviral activity, we analyzed the SEC-purified oligomeric complex by cryo-electron microscopy (cryo-EM) and determined its structure at 2.65 Å resolution (Figure S2 and Table S1). DRT4 assembles into a two-layered hexamer organized as a dimer of trimers, with each trimer forming an approximately equilateral triangle with sides of ∼115 Å (Figure 2A). Consistent with our SEC observations, clear densities corresponding to ssDNA were observed within the complex, allowing five purine nucleotides to be assigned in the channel (Figure 2A, B). The complex contains six DRT4 protomers and six ssDNA molecules, corresponding to a 6:6 stoichiometry, with one ssDNA molecule threaded between the two structural lobes of each protomer (Figure 2A, B). Each DRT4 protomer adopts a bilobed architecture. The N-terminal RT lobe (residues 1–286) comprises the canonical fingers (residues 1–105 and 156–210) and palm (residues 106–155 and 211–286) subdomains conserved across RT families. Unlike canonical RTs^23^, DRT4 lacks a thumb domain. Instead, residues 287–540 fold into a predominantly helical domain that constitutes the C-terminal lobe and is structurally analogous to the corresponding domains in the AbiA, AbiK, and Abi-P2 systems (Figure 2B)^19,20^. Structural alignment revealed that DRT4 most closely resembles AbiK, with a Cα root-mean-square deviation (RMSD) of ∼3.6 Å (Figure S3A). One notable difference is that a loop connecting the fingers and palm domains is substantially longer in DRT4 than in AbiK (Figure S3A). Continuous cryo-EM density connects the 5′ end of the ssDNA to residue Y125 within this extended loop, suggesting that Y125 serves as the priming residue for protein-primed DNA synthesis (Figure 2C). Consistent with this hypothesis, DRT4 displayed a dNTP-dependent mobility shift on SDS–polyacrylamide gel electrophoresis (SDS–PAGE) (Figure 2D), reveal a covalent linkage between the ssDNA product and Y125 and thus a protein-primed mode of ssDNA synthesis, consistent with recent reports^24,25^.

**Figure 2.**
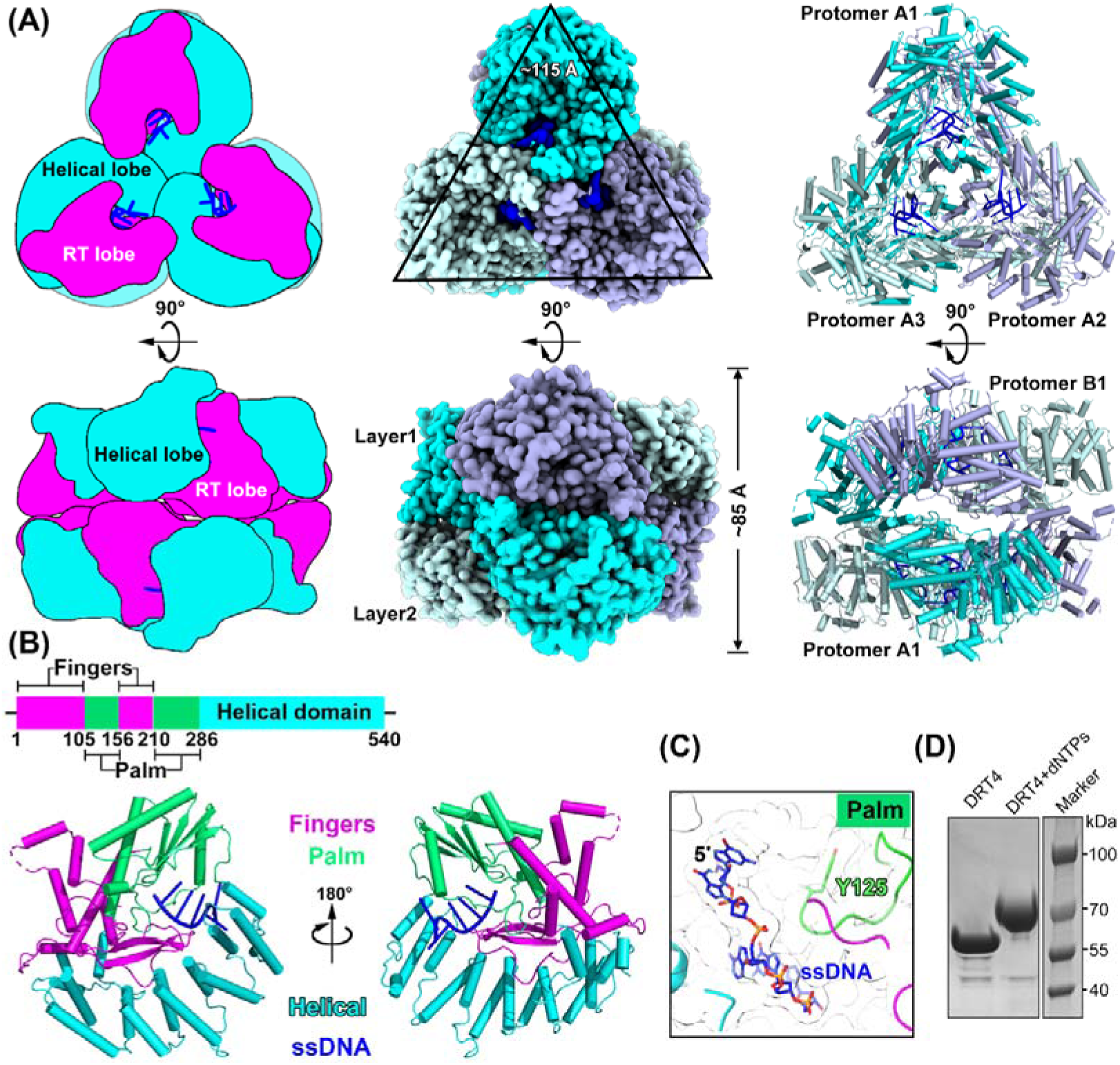
| Overall architecture of the DRT4-ssDNA hexamer. **(A)** Schematic (left), surface (middle), and ribbon (right) representations of the cryo-EM structure of the DRT4–ssDNA complex. **(B)** Domain organization (top) and ribbon representation (bottom) of a DRT4–ssDNA protomer. **(C)** Close-up view showing the 5′ end of the ssDNA covalently linked to residue Y125. **(D)** SDS–PAGE analysis showing that the nascent ssDNA product remains covalently linked to DRT4.

The DRT4–ssDNA hexamer is further stabilized by extensive inter-protomer contacts. Within each trimer, the C-terminal helical lobes assemble into a three-bladed propeller-like arrangement that buries ∼3,600 Å² of surface area and is stabilized by an extensive network of inter-protomer hydrogen-bonding and polar interactions involving residues L136, L287, N287, K300, N360, D320, Q324, and L307 (Figure S3B). In contrast, the interfaces between the two trimeric layers are mediated predominantly by the N-terminal RT lobes. Vertical interfaces between opposing protomers (A1–B1 and equivalents) are connected by salt-bridge and polar interactions involving K173, H204, D466, and E202, whereas diagonal interfaces (A1–B2 and equivalents) are supported by contacts involving T248 and D250 (Figure S3B). Alanine substitution of key interfacial residues (Q320/D324, K173, E202/H204) abolished antiviral activity (Figure S3C), highlighting the essential role of the hexameric assembly in DRT4-mediated defense. Together, these structural features reveal that DRT4 forms a hexameric assembly in which each protomer initiates protein-primed ssDNA synthesis and remains covalently linked to its ssDNA product.

### DRT4 preferentially incorporates purine nucleotides during ssDNA synthesis

The presence of protein-associated ssDNA within the DRT4 hexamer suggested that DRT4 actively synthesizes DNA. To test whether purified DRT4 could extend these DNA products *in vitro*, we incubated DRT4 with dNTPs in the presence of various divalent metal ions. At physiologically relevant concentrations, only Mg² supported DRT4-mediated DNA synthesis, yielding short products of <100 nt. Increasing the dNTP concentration above 50 µM did not further increase either the length or abundance of the synthesized DNA products (Figure 3A and Figure S3D). To determine the nature of these products, the reaction mixtures were treated with either nuclease P1, which preferentially degrades ssDNA, or Duplex DNase, which selectively degrades dsDNA. SDS–PAGE analysis showed that nuclease P1 treatment eliminated the slowly migrating DRT4–DNA species and restored the migration of free DRT4, whereas Duplex DNase had no obvious effect (Figure 3B). These results indicate that DRT4 synthesizes protein-linked ssDNA in a template-independent manner.

**Figure 3.**
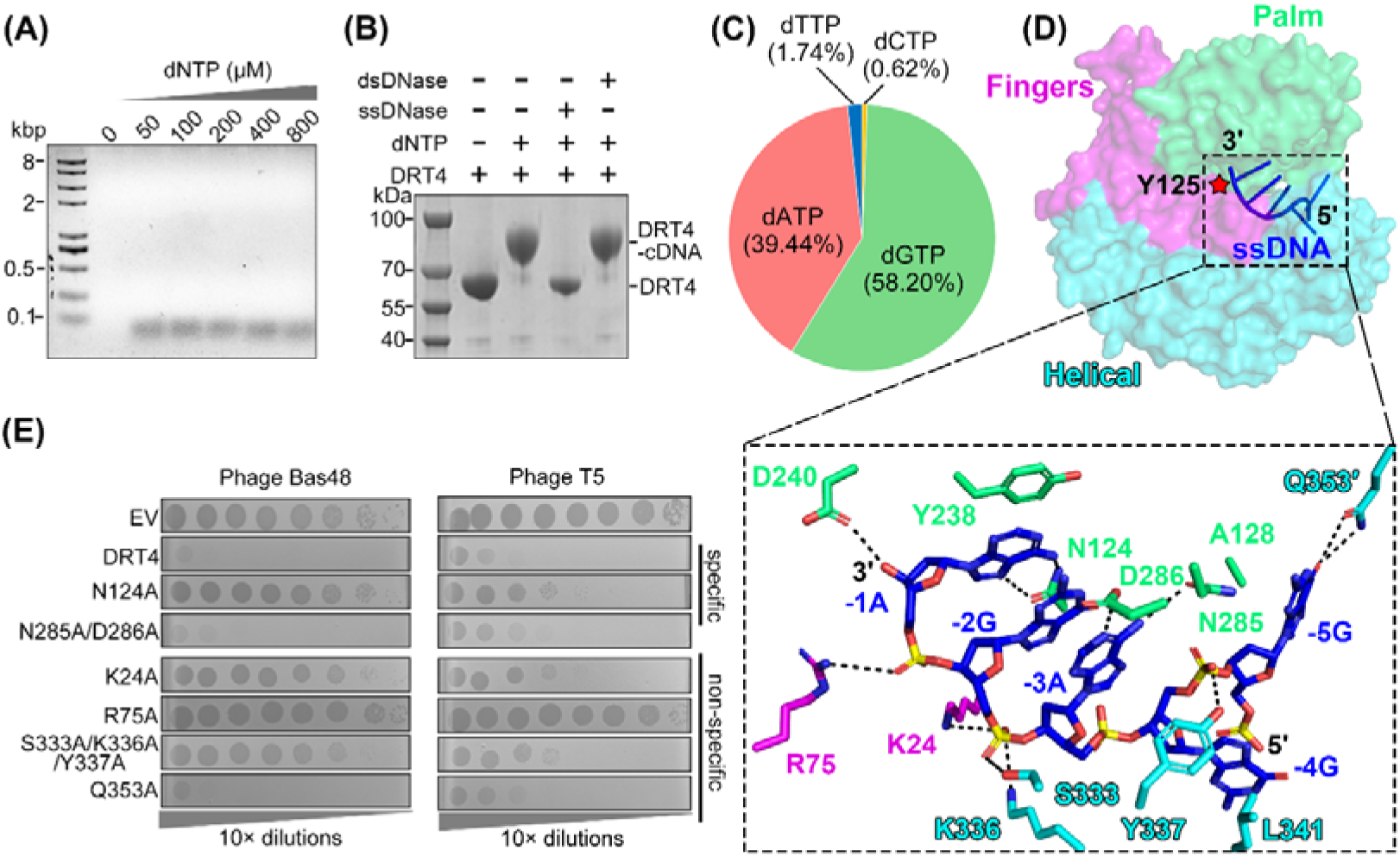
| DRT4 preferentially synthesizes purine-rich ssDNA. **(A)** Gel analysis of DNA products synthesized by DRT4 using increasing concentrations of mixed dNTPs in the presence of 5 mM Mg² . DRT4 (400 nM) was incubated with 0, 50, 100, 200, 400, or 800 μM dNTPs. **(B)** Nuclease sensitivity analysis of DRT4-synthesized DNA products. Following incubation of DRT4 with 800 μM dNTPs at 37°C for 15 min, reaction products were treated with nuclease P1 or Duplex DNase at 37°C for 30 min. **(C)** Mass spectrometric analysis of the nucleotide composition of DNA synthesized *in vitro* by the DRT4–ssDNA complex. **(D)** Overall (top) and close-up (bottom) views of the ssDNA synthesis channel in DRT4. **(E)** Effects of mutations in the ssDNA synthesis channel on DRT4-mediated antiphage activity. Images are representative of three biological replicates.

To define the nucleotide composition of the DRT4-generated DNA, we isolated the *in vitro* synthesized products and analyzed them by mass spectrometry. DRT4 displayed a striking preference for purine incorporation, with dGTP and dATP accounting for 58.20% and 39.44% of incorporated nucleotides, respectively, whereas dTTP and dCTP represented only 1.74% and 0.62% (Figure 3C). Structural analysis provided a potential basis for this purine selectivity. At the nucleotide-binding site, N124 forms bidentate hydrogen bonds with the N6 and N7 atoms of adenine or probably the O6 and N7 atoms of guanine, thereby favoring recognition of purine bases. N285 and D286 further contribute to purine recognition of the nucleotide positioned at −3 through additional base-specific interactions (Figure 3D). Mutations of residues involved in nucleotide recognition (N124A, N285A/D286A) impaired anti-phage activity (Figure 3E), indicating a functional role for purine-selective DNA synthesis in DRT4-mediated defense.

The structure further revealed how the nascent ssDNA is stabilized within the DRT4 complex. The nucleotides at positions −1 to −3 relative to the catalytic center adopt an extended, side-by-side configuration and are stabilized predominantly through interactions with the phosphate backbone contributed by both the RT and C-terminal helical lobes. Residues K24, R75, and D240 from the fingers and palm subdomains, together with S333 and K336 from the helical domain, participate in these contacts. At position −4, the phosphate group is coordinated by Y337, promoting a characteristic flipped-out conformation that is further stabilized by L341. The nucleobase at position −5 forms additional contacts with A128 from the same protomer and Q353 from an adjacent protomer, thereby extending the DNA-binding interface across neighboring subunits (Figure 3D). Consistently, mutating representative residues (K24A, R75A, S333A/K336A/Y337A, Q353A) involved in these nucleic acid contacts reduced DRT4-mediated anti-phage defense (Figure 3E). Together, these findings show that DRT4 catalyzes template-independent synthesis of protein-linked ssDNA with a pronounced preference for purine nucleotides, while an extensive DNA-binding network spanning the RT and C-terminal helical domains stabilizes the synthesized DNA and is required for antiviral defense.

### Phage 0087 and NrdAB activate DRT4 to trigger cell death

We next sought to identify the phage-encoded factors that trigger DRT4 activation. Infection of DRT4-expressing *E. coli* with phages Bas48 and Bas49 resulted in strong protection with rare phage escapers emerged. We reasoned that these escapers might harbor spontaneous mutations that enable evasion of DRT4-mediated defense. To identify the genetic basis of escape, we isolated multiple independent escapers from each phage and subjected them to whole-genome sequencing. Bas48 escapers carried mutations in the gene encoding hypothetical protein 0087 (H260Q or a P232+C frameshift), or a mutation in hypothetical protein 0082 (D97E) accompanied by a mutation in the ribonucleotide reductase subunit NrdA (Y213C). Similarly, Bas49 escapers exclusively harbored mutations in the gene encoding the Bas49 NrdA homolog (R209C or S485L) (Figure 4A).

**Figure 4.**
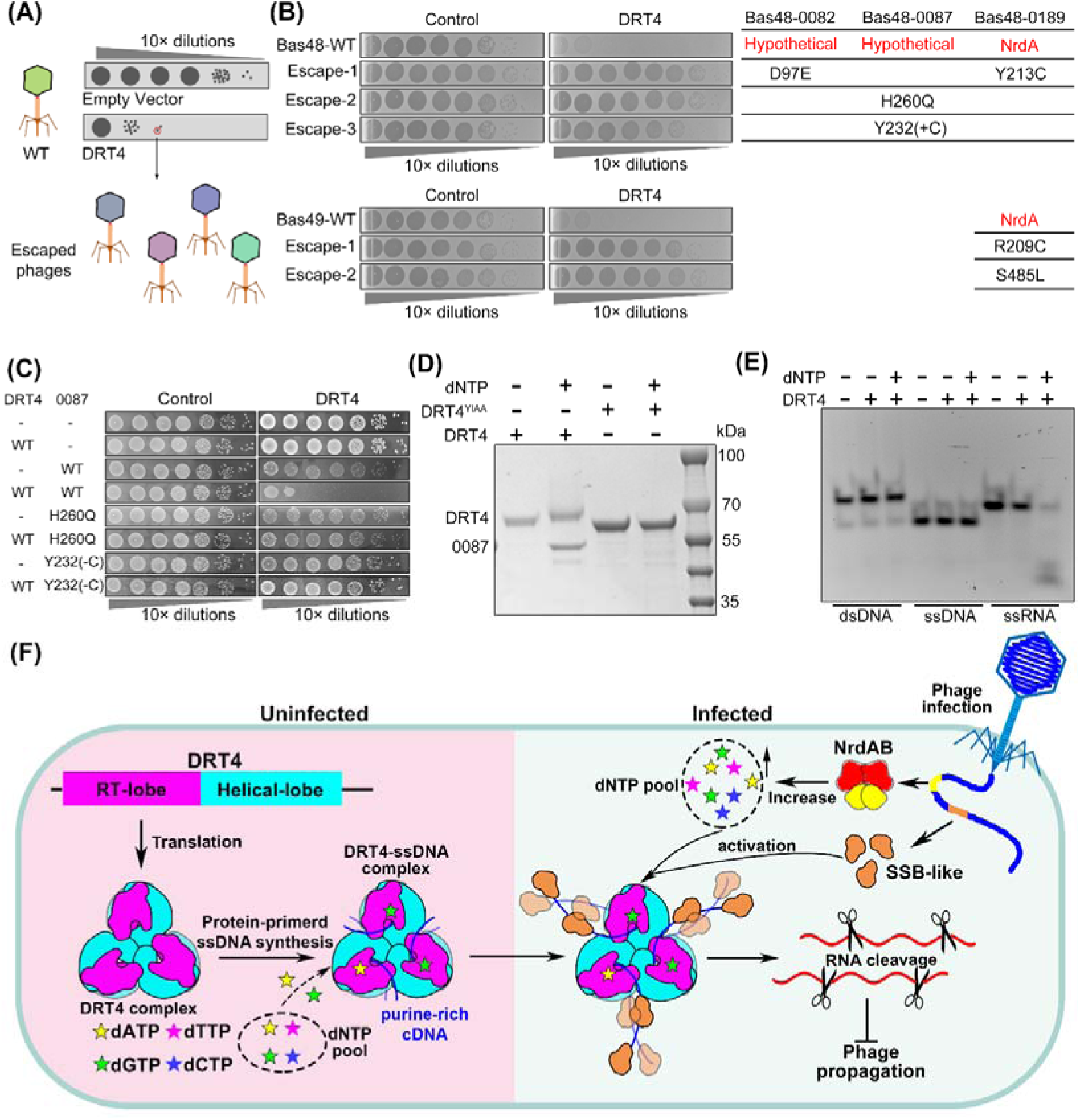
| Phage SSB-like protein and dNTP activate the DRT4 immunity. **(A)** Plaque assays of isolated phage escapers against the DRT4 defense system. Serial dilutions of the indicated phages were spotted onto bacterial lawns expressing DRT4. Escape-associated mutations are indicated on the right. **(B)** Growth or survival of *E. coli* cells co-expressing DRT4 with wild-type or mutant Bas48 NrdAB. Images are representative of three biological replicates. **(C)** Growth or survival of *E. coli* cells co-expressing DRT4 with wild-type or mutant Bas48 0087. Images are representative of three biological replicates. **(D)** *In vitro* pull-down assay showing that Bas48 0087 associates with the ssDNA product synthesized by DRT4. **(E)** Cleavage assays using ssDNA, dsDNA, or ssRNA substrates in the presence or absence of dNTPs. **(F)** Proposed model for DRT4-mediated antiphage immunity.

Given the high sequence similarity between the corresponding proteins of Bas48 and Bas49, we focused on Bas48 to determine whether the candidate phage proteins could directly activate DRT4. Phage-encoded ribonucleotide reductases (RNRs), typically composed of the NrdA and NrdB subunits, catalyze the conversion of ribonucleotides into deoxyribonucleotides, thereby supporting the dNTP supply required for phage DNA replication^26,27^. To test whether the candidate phage factors could trigger DRT4-dependent toxicity, we expressed Bas48 0087 or co-expressed nrdA and nrdB in *E. coli* cells with or without DRT4 and monitored bacterial growth. Expression of either 0087 or nrdAB markedly inhibited growth specifically in DRT4-containing cells, whereas little effect was observed in cells lacking DRT4 (Figure 4B, C). In contrast, the escape-associated 0087 variants failed to induce DRT4-dependent growth inhibition (Figure 4C). These results identify phage protein 0087 and NrdAB as independent phage-encoded activators of the DRT4 defense system.

### The SSB-like protein 0087 directly binds the synthesized ssDNA product

To investigate how phage 0087 and NrdAB activate the DRT4 system, we first examined the predicted structure of 0087. The protein comprises three structured domains connected by flexible linkers, followed by a C-terminal acidic tail (Figure S4A). Domain II adopts a canonical OB fold and shows structural similarity to prokaryotic single-stranded DNA-binding proteins (SSBs)^28^ and eukaryotic replication protein A (RPA) (Figure S4B). Moreover, the C-terminal tail terminates in a conserved “DDIPF” motif resembling the C-terminal tip of *E. coli* SSB, a region known to mediate interactions with partner proteins (Figure S4C)^29^. Given our previous finding that phage SSB proteins activate DRT2, together with reports that SSB-like proteins recognize oligoadenylate products generated by DRT9^9^, we hypothesized that 0087 might activate DRT4 by binding its ssDNA product.

To test this hypothesis, we performed *in vitro* pull-down assays. 0087 specifically co-precipitated with wild-type DRT4 in the presence of dNTPs, whereas no detectable interaction was observed with the YIAA mutant, which is defective in ssDNA synthesis (Figure 4D). These results indicate that the interaction between 0087 and DRT4 depends on DRT4-mediated DNA synthesis, consistent with 0087 directly binding to the newly synthesized ssDNA product. This finding is consistent with a recent report showing that T7 SSB promotes the RNA cleavage activity of the DRT4 system^24^, suggesting that SSB activates DRT4-mediated immunity not by promoting DNA synthesis but by inducing a conformational change in ssDNA. A BLASTp search identified close homologs of 0087 in several related phages, including Bas49, Bas50, and Bas52–59 (Figure S5A). DRT4 conferred strong protection against a subset of these phages, including Bas49 and Bas50, reducing EOP by up to ∼10^6^-fold (Figure 1B). Similarly, a recent study reported that the 0087 protein from the Bas52 system directly activates the DRT10 system^14^. Collectively, these findings suggest that 0087-like SSB proteins represent a conserved class of phage-encoded factors capable of binding ssDNA generated by diverse DRT systems and triggering their subsequent activation and defense.

### Activated DRT4 cleaves RNA in a dNTP-dependent manner

Our results indicate that DRT4 can be activated through at least two phage-associated signals: direct recognition of the DRT4-generated ssDNA product by the SSB-like protein 0087 and perturbation of nucleotide metabolism associated with phage NrdAB. The latter is reminiscent of our previously characterized DRT2 and DRT9 systems^8,9^, in which phage-encoded NrdAB elevates intracellular dNTP levels to promote defense activation, whereas phage escape mutations in NrdAB impair ribonucleotide reduction and thereby evade immunity. These parallels suggest that sensing phage-induced changes in nucleotide metabolism may represent a shared activation strategy among diverse DRT systems. We next sought to determine how activated DRT4 executes its antiviral function. Previous work showed that DRT7 possesses nuclease activity in addition to its reverse-transcriptase activity^30^, prompting us to test whether DRT4 similarly exhibits nucleolytic activity. Purified DRT4 was incubated with FAM-labeled ssDNA, dsDNA, or ssRNA substrates in the presence or absence of dNTPs. DRT4 readily cleaved ssRNA in the presence of dNTPs, whereas no obvious cleavage of ssDNA or dsDNA was detected under the tested conditions (Figure 4E). Addition of 0087 did not further enhance ssRNA cleavage in the presence of dNTPs (Figure S5B). This observation is consistent with recent studies proposing that phage SSB proteins promote DRT4 activation by binding and displacing the newly synthesized ssDNA from the polymerase pocket, whereas dGTP binding is additionally required to establish the RNase-active state^24^. Because our *in vitro* reactions contained all four dNTPs, including dGTP, DRT4 may already have been driven toward an RNase-active state, thereby limiting any additional stimulatory effect of 0087 under these conditions. *In vivo*, 0087 may instead promote activation by binding and stabilizing the DRT4-generated ssDNA from the cellular nuclease and facilitating its displacement from the catalytic pocket. Together, these results indicate that DRT4 possesses a dNTP-dependent RNase activity, suggesting RNA cleavage as the downstream effector mechanism of activated DRT4. This is consistent with a recent report showing that activated DRT4 removes 3′-terminal guanosine monophosphate residues from both phage and host RNAs, thereby terminating infection^24^.

## Discussion

In this study, we identify two distinct phage-derived signals that activate the DRT4 immune system and connect phage sensing to RNA cleavage and abortive infection. DRT4 assembles into a two-layered homohexamer that catalyzes protein-primed, template-independent ssDNA synthesis with a strong preference for purine nucleotides. Upon phage infection, DRT4 can be activated through two distinct viral cues: direct binding of the phage-encoded SSB-like protein 0087 to the newly synthesized ssDNA or increased in intracellular dNTP levels associated with the phage-encoded NrdAB ribonucleotide reductase complex. Activated DRT4 subsequently cleaves RNA, leading to host cell death through abortive infection and thereby restricting phage propagation (Figure 5F).

DRT4 therefore expands the functional repertoire of defense-associated reverse transcriptases beyond canonical RNA-templated reverse transcription. Consistent with recent studies^24,25^, DRT4 possesses protein-primed DNA polymerase and RNA endonuclease activities, with ssDNA synthesis and phage DNA-binding proteins contributing to activation of its RNA-cleaving effector function. Rong et al. further proposed that displacement of the synthesized ssDNA from the polymerase pocket promotes a conformational rearrangement of the DRT4 hexamer, leading to formation of an RNA-binding and cleavage site^24^. Our results complement this model by identifying physiological phage-encoded triggers through escape genetics. During phage infection, NrdAB-driven changes in intracellular dNTP levels promote the synthesis of the DRT4-linked ssDNA, whereas the phage SSB-like protein 0087 directly binds the newly synthesized ssDNA and facilitates conversion of DRT4 into an RNase-active state. Activated DRT4 subsequently cleaves host and phage RNA, thereby restricting phage replication and triggering abortive infection^24^. Thus, DRT4 uses its own DNA product as a signaling intermediate to integrate phage-induced metabolic changes and direct signals of active viral replication into an antiviral response.

Dual sensing of phage infection also appears to be a common feature of DRT-mediated immunity. In our recent study of DRT2^8^, the system responds to both NrdAB-associated changes in dGTP level within nucleotide pools and phage-encoded single-stranded DNA-binding proteins (SSBs) or single-strand annealing proteins (SSAPs) produced during active viral replication. DRT2 first synthesizes an ssDNA intermediate, suggesting that the newly generated DNA may serve as a molecular sensor or scaffold for detecting phage replication proteins. Similarly, our previous work on DRT9 showed that elevated dATP levels associated with phage NrdAB promote synthesis of the antiviral ssDNA product^9^. Because newly synthesized DNA is continuously counteracted by cellular nucleases, increased dNTP availability during phage infection may shift the balance toward ssDNA accumulation for anti-phage defense. Phage-encoded SSBs and SSAPs are central components of viral DNA replication^31,32^. Increasing evidence suggests that these proteins can also be sensed through RT-generated ssDNA intermediates, where their binding stabilizes or remodels the DNA and promotes downstream immune activation. Mutations in these proteins can enable escape from diverse defense systems, including AbiK^21^, DRT2^8^, Retron-Eco8, Hachiman, ShosA, and Borvo^33^. Notably, such interactions appear particularly common among RT-associated defense systems. For example, Retron-Eco8 detects T7 SSB through direct binding of its msDNA, and disruption of the ssDNA-binding activity of SSB abolishes Retron-Eco8 activation^34–36^. Likewise, in the mechanistically related DRT2 and AbiK system, phage escape can occur through mutations in SSAP proteins^8,21^. Together with our findings for DRT4, these observations suggest a broader principle in which RT-generated ssDNA serves as an infection-sensing platform that integrates phage-induced metabolic changes with direct recognition of essential viral replication proteins.

Template-independent nucleic acid synthesis has been documented in several classes of ribonucleotidyl transferases (rNTrs), which catalyze the addition of one or more ribonucleoside triphosphates to the 3′ hydroxyl group of a primer^37,38^. While most rNTrs exhibit specificity for particular ribonucleotides, CCA-adding enzymes specifically incorporate the trinucleotide CCA at the 3′ end of tRNA precursors. A related principle is observed in RT-like defense proteins in which the canonical thumb domain is replaced by a helical domain, including AbiA^20^, AbiK^17^, and Abi-P2^19^, where template-independent synthesis of ssDNA from deoxynucleotides appears to be a conserved feature^19,20^. Although AbiK can utilize all four deoxynucleotides with relatively limited bias, other members of this family display pronounced nucleotide preferences. For example, AbiA preferentially incorporates adenine and cytosine^20^. Recent studies of DRT3 and DRT7 further suggest that amino-acid residues within these enzymes can directly influence nucleotide selection during template-independent DNA synthesis, thereby allowing the protein itself to encode sequence bias in the synthesized DNA product^11,12,39,40^. In DRT4, our structural and biochemical analyses reveal that nucleotide selectivity is governed by specific interactions between purine bases and residues within the catalytic pocket. These observations suggest that direct base recognition by the polymerase active site may represent a broader mechanistic principle among helical-domain RT defense systems. Such protein-directed nucleotide selection could enable different systems to generate compositionally distinct ssDNA products, which may in turn acquire specialized structural or signaling properties during antiviral defense.

In summary, our study, together with two recent parallel studies^24,25^, expands our understanding of how DRT4 mediates antiphage defense. our study reveal how DRT4 integrates phage-induced nucleotide changes and direct recognition of a phage SSB-like protein to activate RNA cleavage and abortive infection. DRT4-generated ssDNA serves as a key signaling intermediate linking phage sensing to antiviral effector activation, further expanding the diverse roles of reverse transcriptases in bacterial immunity.

## MATERIALS AND METHODS

### Plasmid and *E. coli* strain construction

*Escherichia coli* DH5α, BL21 Star (DE3) and MG1655 cells were used for plasmid construction, protein expression and phage related experiments, respectively. Phages T4, T5 and T7 were gifts from Xiao Yi’s lab at Shenzhen Institutes of Advanced Technology, Chinese Academy of Sciences. Phages T1(11303-B1), T2 (11303-B2) and T6 (11303-B6) were obtained from the American Type Culture Collection (ATCC). Phage lambda (DSM 4499) and isolates from the·BASEL collection were acquired from the Leibniz Institute DSMZ-German Collection of Microorganisms and Cell Cultures.

### Plasmid cloning, protein expression and purification

The *Escherichia coli* 21-C8-A *drt4* gene, including a 200-bp upstream region and a C-terminal Strep-tag, was synthesized by Sangon Biotech and cloned into the pRSFDuet-1 expression vector. The resulting construct (pRSFDuet-DRT4-Strep) was introduced into *E. coli* BL21 Star (DE3). Protein expression was induced by the addition of 0.5 mM isopropyl β-D-1-thiogalactopyranoside (IPTG; Sangon Biotech) when the culture reached an OD_600_ of approximately 0.8. Following induction, the cultures were incubated at 18 for 20 h. Cells were harvested by centrifugation and resuspended in lysis buffer containing 20 mM Tris-HCl (pH 8.0), 300 mM NaCl, and 2 mM dithiothreitol (DTT).

For DRT4 purification, cell pellets were disrupted and lysed by AH-1500 High Pressure Homogeniser (ATS, inc.) at 800 bar for 10 min and centrifuged for 30 min at 32,914 × g. The supernatant was loaded onto a Strep-tag II affinity column (Smart Lifesciences). The column was washed five times with lysis buffer, and the bound proteins were eluted using buffer A (20 mM Tris-HCl, pH 8.0, 100 mM NaCl, and 1 mM DTT) supplemented with 20 mM D-biotin. The eluted protein fractions were subsequently loaded onto a 5 mL HiTrap Q Fast Flow column (Cytiva) and separated using a linear NaCl gradient ranging from 100 mM to 2 M over 20 column volumes. The fractions containing DRT4 were further purified by size-exclusion chromatography using a Superdex 200 Increase 10/300 GL column (Cytiva) equilibrated with buffer containing 20 mM Tris-HCl (pH 8.0), 300 mM NaCl, and 2 mM DTT. All DRT4 mutants were generated by site-directed mutagenesis and purified using the same procedure as described for the wild-type protein.

### Phage plaque assays

The *Escherichia coli* 21-C8-A DRT4, along with its upstream 200 bp sequence was cloned into the pACYC184 plasmid. The resulting pACYC184-DRT4 plasmid and the empty pACYC184 vector (EV) were separately transformed into *E. coli* MG1655. Transformants were selected on LB agar plates containing 10 μg/mL tetracycline at 37 . Single colonies were inoculated into liquid LB medium supplemented with the appropriate antibiotic and cultured overnight at 37 with shaking at 200 rpm. On the following day, 500 μL of the overnight culture was mixed with 25 mL of molten MMB top agar (LB supplemented with 0.1 mM MnCl , 5 mM MgCl , 5 mM CaCl , 0.5% agar) and poured onto square plates (Biosharp). The plates were allowed to dry for 1 hour at room temperature. Tenfold serial dilutions of phage suspensions were prepared in phosphate-buffered saline (PBS), and 10 μL of each dilution was spotted onto the corresponding bacterial lawns. The plates were incubated overnight at 37 , after which plaques were counted to determine the phage titer, expressed as plaque-forming units per milliliter (PFU/mL)

### Quantification of dNTPs in cDNA products by UPLC-MS

The cDNA products were performed by incubating 400 nM of DRT4 with 800 µM dNTP mixtures and 5 mM MgCl_2_ at 37°C for 120 min. cDNA was extracted as previously described. The purified cDNA products were subjected to enzymatic hydrolysis using Exonuclease I (Exo I) for 1 h. Following incubation, the reaction mixtures were centrifuged at 13,000 rpm for 10 min, and the supernatant was used for UPLC-MS analysis.

### *In vivo* growth toxicity assays

Wild-type or mutant variants of plasmids pACYC184-DRT4 and pBAD-Bas48-0087 pBAD-Bas48-NrdAB were co-transformed into *E. coli* Mg1655 competent cells. Transformed colonies were cultured overnight at 37°C in LB medium supplemented with 2% glucose and dual antibiotics (10 μg/mL Tetracycline and 100 μg/mL ampicillin). A fresh LB-glucose medium containing identical antibiotics was inoculated with 1% (v/v) of the overnight culture and incubated at 37°C with shaking until mid-log phase growth (OD600= 0.6). Cells were harvested by centrifugation (4,500 rpm, 2 min) and subjected to two sequential washes using antibiotic-free, glucose-free LB medium, with each wash step performed under identical centrifugation conditions (4,500 rpm, 2 min). The washed cell pellets were resuspended in LB medium containing 2% glucose(m/v) for inhibiting proteins expression or 0.02% L-arabinose(m/v) and incubated at 37 °C (250 rpm) for 4H. Then serially diluted through ten-fold increments using antibiotic-free, glucose-free LB medium. 10 µL aliquots were then taken from each dilution and pipetted onto LB agar plates supplemented with the additives (2% glucose for inhibiting proteins expression or 0.02% L-arabinose for induced expression) and the same antibiotics as mentioned. The plates were subsequently incubated at 37 °C overnight.

### *In vitro* cDNA synthesis assays

Reactions were carried out by adding indicated concentrations of DRT4 protein or with Bas48-087 protein, then incubated with 800 μM deoxynucleotides into a reaction buffer containing 20 mM Tris-HCl (pH 8.0), 150 mM NaCl, and 5 mM MgCl_2_. The final reaction volume was 100 μL, and reactions were incubated at 37°C for 180 minutes unless otherwise specified. To remove the DRT4 protein, 4 mg/mL proteinase K (TIANGEN) was added, and the solution was incubated at 65°C for 30 minutes. To extract the cDNA, the protease-treated solution was heated to 95°C for 10 minutes to inactivate proteinase K. The cDNA was precipitated by adding an equal volume of pre-cooled isopropanol and incubating for 2 hours at -80°C, followed by centrifugation at 21,130 g for 15 minutes. The supernatant was discarded, and the pellet was washed once with 75% ethanol solution and allowed to dry. The final cDNA samples were dissolved in 50 μL of ddH_2_O. Then, the cDNA products were analyzed by running 1% agarose gels in 1× tris-acetate-EDTA buffer, followed by staining with SYBRGOLD (Invitrogen, S11494) and visualizing with a ChempGel 7000 molecular imager (SageCreation Beijing, China)

### Pull down assay

Purified Strep-tagged DRT4 were incubated with 800 μM dNTPs in a binding buffer (20 mM Tris–HCl, pH 8.0, 150 mM NaCl, 5mM Mg^2+^) at 37°C for 15 min. His-tagged Bas48-087 at 4°C for 3 hours in the presence of 800 μM dNTPs in a binding buffer (20 mM Tris–HCl, pH 8.0, 150 mM NaCl, 5mM Mg^2+^). The complex was loaded onto a Strep-tag II column (Smart Lifesciences), washed three times with binding buffer, and proteins were eluted with binding buffer containing 20 mM D-biotin. Eluted proteins were analyzed by 4-20% SDS-PAGE.

### *In vitro* protein shift assays

Reactions were carried out by adding 1 mg/mL DRT4 protein and 800 μM deoxynucleotides into a reaction buffer containing 20 mM Tris-HCl (pH 8.0), 150 mM NaCl, and 5 mM MgCl_2_, reactions were incubated at 37°C for 30min, then ssDNase (NEB, M0502) or dsDNase (Beyotime, D7028S) were added to the mixture. Samples were analyzed by 4-20% SDS-PAGE.

### dsDNase、ssDNase and RNase activity assay

Cleavage reactions were conducted in 10 μL reaction buffer (50 mM Tris-HCl pH 7.5, 75 mM KCl, 1 mM MgCl ) containing 200 nM 5′-6-FAM-labeled dsDNA, ssDNA, ssRNA substrate. Reactions were initiated by adding 1 μM of either DRT4 alone, DRT4 with Bas48-087 protein, proceeded at 25°C for 30 minutes. To remove protein, 4 mg/mL proteinase K (TIANGEN) was added, and the solution was incubated at 65°C for 30 minutes. After incubating at 95°C for 5 minutes, the reaction mixtures were electrophoresed on 12% denaturing polyacrylamide gels and visualized using a Tanon 6100 Multi imaging system (Tanon Science & Technology Co., Ltd., Shanghai, China). The cleavage assays were repeated at least three times, and representative results are shown.

### Cryo-EM sample preparation and data acquisition

3.5 μL of a ∼11 mg/mL solution of purified DRT4-ssDNA complex is applied onto glow-discharged UltrAuFoil 300 mesh R1.2/1.3 grids (Quantifoil). The grids were blotted for 2 s at 100% humidity and flash frozen in liquid ethane using a FEI Vitrobot Mark IV (FEI). Images were collected on a Titan Krios electron microscope (FEI) operated at an acceleration voltage of 300 kV, equipped with a Falcon4 detector with a physical pixel size of 0.838 Å. The defocus during image collection ranged from -1.5 µm to -2.5 µm. Images were dose-fractionated into 32 frames with a total accumulated dose of 50 electrons per Å^2^.

### Cryo-EM data processing

Image processing was conducted by RELION 3.1^41^ and cryoSPARC v4.3^42^. Motion correction was performed with MotionCor2^43^. Contrast transfer function (CTF) parameters were estimated by Ctffind4^44^. Auto-picked particles using the Laplacian-of-Gaussian method were extracted and underwent two rounds of 2D classification to remove junk particles and several rounds of heterogeneous refinement in cryoSPARC v4.3. A total of 824,252 particles were used for Ab-Initio reconstruction of DRT4 complex. After iterations of heterogeneous refinement, particles corresponding to the best class with the highest-resolution features were selected and subjected to non-uniform refinement in cryoSPARC v4.3. These particles then underwent CTF refinement followed by another round of non-uniform refinement, generating the final reconstitutions. All resolutions were estimated by applying a soft mask around the protein density and the Fourier shell correlation (FSC) = 0.143 criterion in cryoSPARC v4.3. Local resolution estimates were calculated from two half data maps in cryoSPARC v4.3. Further details related to data processing and refinement are summarized in Supplementary Table 1.

### Model building and refinement

DRT4 atomic model was built with the help of AF3 predicted structure, and then manually refined by COOT v0.9.5^45^ interactively. All models were then refined against combined maps using phenix.real_space_refine v1.19.2^46^ by applying geometric and secondary structure restraints. All structural figures were prepared in Pymol v2.4.2 (http://www.pymol.org) and Chimera X^47^.

## ACKNOWLEDGEMENTS

We thank the staff at Southern University of Science and Technology (SUSTech) Cryo EM Center for assistance in data collection on the SUSTech Titan KRIOS cryo-electron microscope. This work was funded by the National Key R&D Program of China (2024YFE0198900); National Natural Science Foundation of China (grant no. 32625011); Shenzhen Natural Science Foundation (grant no. QNXMA20250701094715021); HE Science Foundation (grant no. HERF2025003).

N.J. is an investigator of SUSTech Institute for Biological Electron Microscopy.

## AUTHOR CONTRIBUTIONS

Xin-Yang Wei, Hua Qi, Xin-Yi Song, Linshu Li: Data curation; formal analysis; methodology. Yushan Xia, Jun-Tao Zhang, Ning Jia,: Data curation; formal analysis; Writing—review and editing. Ning Jia: Project administration.

## ETHICS STATEMENT

This study involved no animal or human experiments. There are no ethical issues involved.

## DATA AVAILABILITY

The cryo-EM density maps have been deposited in the Electron Microscopy Data Bank (EMD-82404 for DRT4-ssDNA complex [https://www.ebi.ac.uk/emdb/EMD-82404]. The atomic coordinates have been deposited in the Protein Data Bank (PDB) with accession number 43YZ (DRT4-ssDNA complex) [https://www.rcsb.org/structure/43YZ]. This paper does not report original code. Source data are provided with this paper.

