## Supplementary material for "Dual phage sensing activates an RNA-targeting reverse transcriptase for anti-phage defense": Figures S1-S5

### Supplementary figures and table

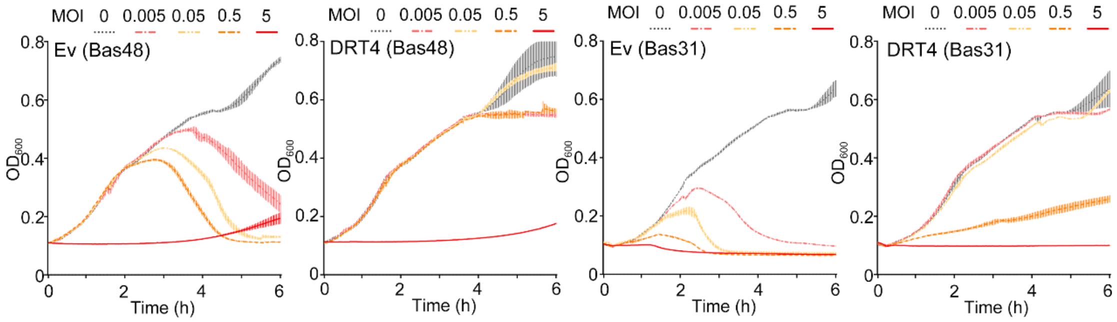


**Figure S1 | DRT4 protects cells through abortive infection.**

Growth curves of *E. coli* strain MG1655 in liquid culture transformed with the DRT4 system or an empty vector (Ev) infected by Bas48 or Bas31 phages. Bacteria were infected at time 0 at an MOI of 0 (no phage), 0.005, 0.05, 0.5, or 5.

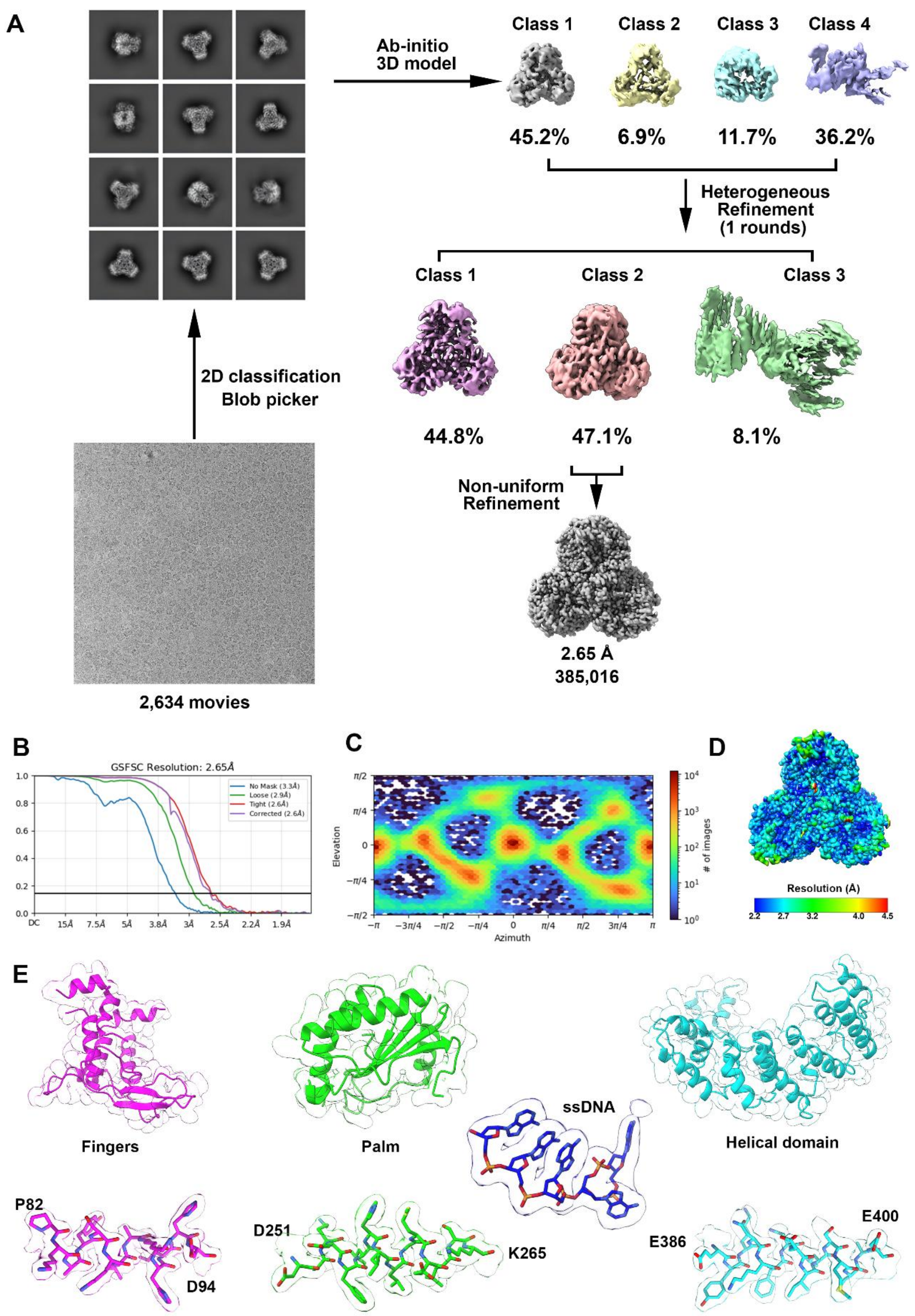


**Figure S2 | Cryo-EM reconstruction of the DRT4-ssDNA complex.**

**(A)** Flow chart of image processing for DRT4-ssDNA complex.

**(B)** Fourier Shell Correlation curve of DRT4-ssDNA complex.

**(C)** Direction distribution plot of DRT4-ssDNA complex.

**(D)** Final 3D reconstructed map of DRT4-ssDNA complex, colored according to local resolution.
**(E)** Cryo-EM densities of each domain of DRT4-ssDNA complex. Densities for the indicated regions are shown in the context of the atomic model.

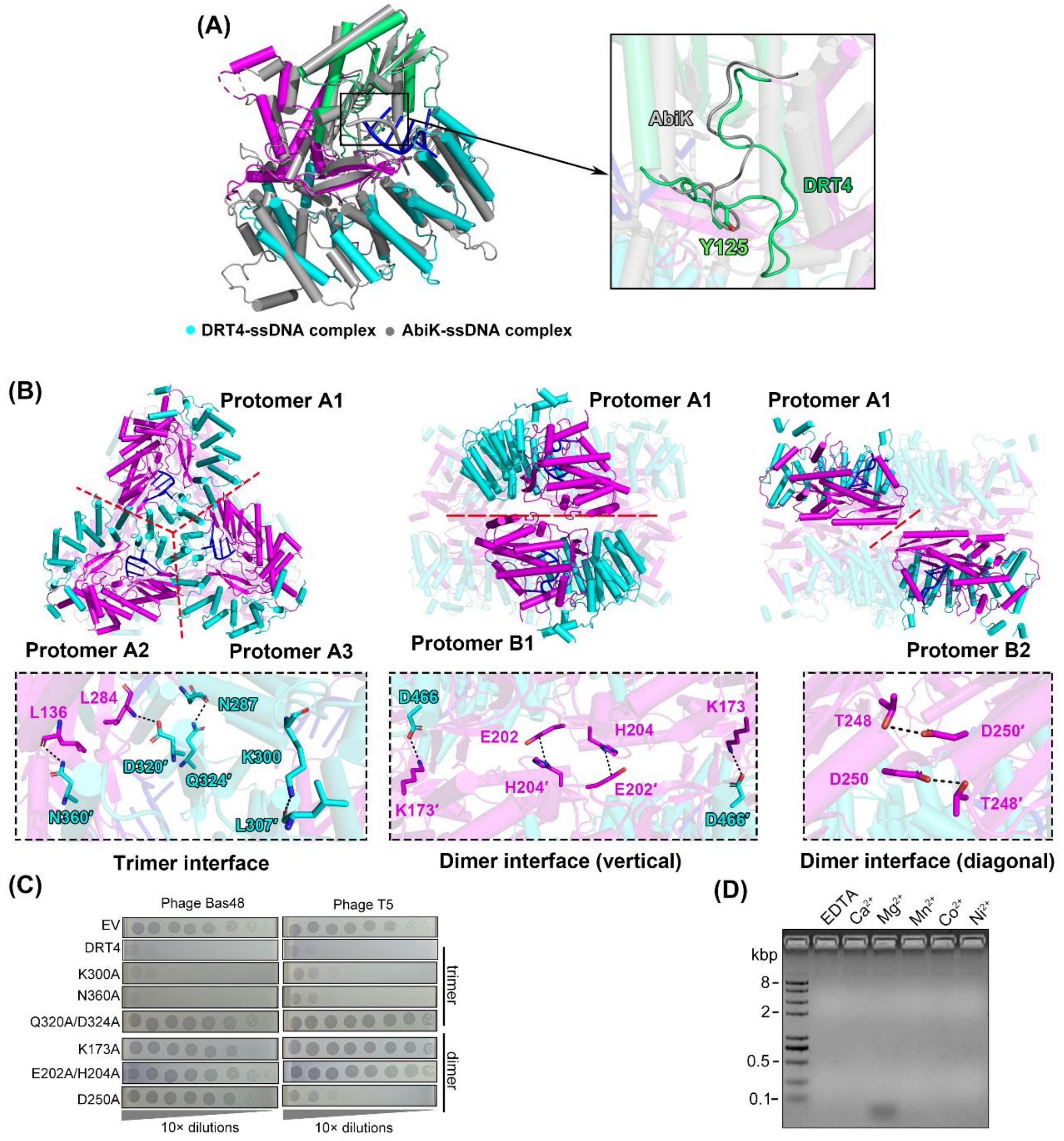


**Figure S3 | Structural comparison of DRT4 with related proteins and the essential role of DRT4–ssDNA complex hexamerization in antiphage activity.**

**(A)** Structural comparison of DRT4-ssDNA complex (in color) with AbiK-ssDNA complex (7R06, in grey).

**(B)** Interactions in the trimer (left) and dimer (middle and right) interfaces contribute to the formation of the DRT4-ssDNA hexamer complex. Detailed interactions are shown in the insets below.

**(C)** Antiphage activity of wild-type DRT4 and interface-related mutants. Images are representative of three replicates.

**(D)** Gel analysis of cDNAs synthesized by DRT4 using different divalent metal ions at physiological concentrations, respectively. Reactions were performed by incubating DRT4 (400 nM) with 800 μM dNTP mixtures.

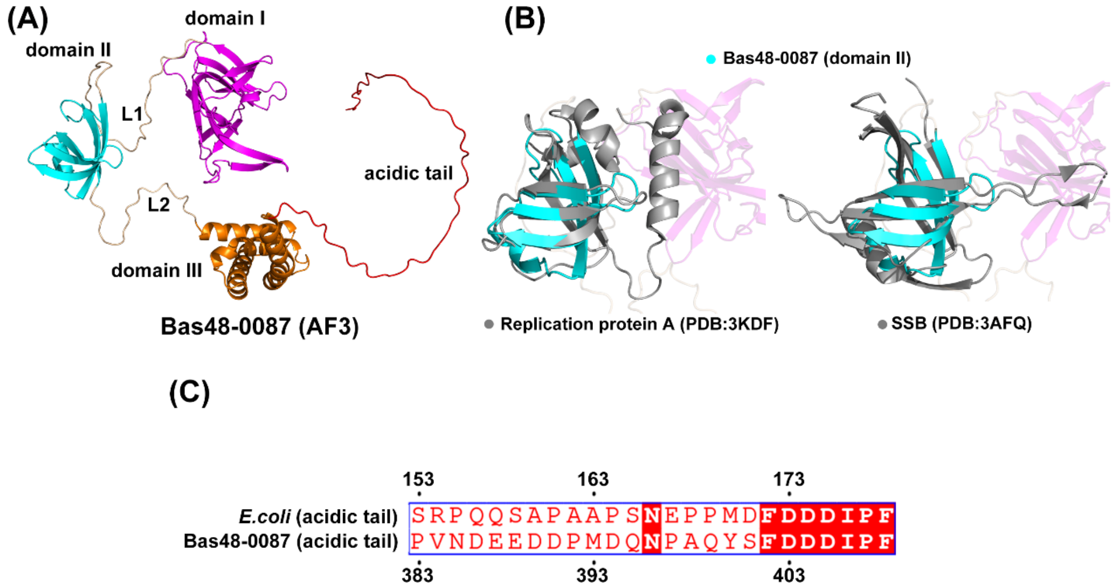


**Figure S4 | Bas48-0087 is an SSB-like protein.**

**(A)** AF3-predicted Bas48-0087 structure, which was colored according to domains.

**(B)** Structural comparison of the domain II of Bas48-0087 (in color) with replication protein A (left, 3KDF, in grey) and single strand binding protein (right, 3AFQ, in grey).

**(C)** Sequence alignment of the acidic tail form the SSB protein of *E.coli* and Bas48-0087 indicated they shared a conserved DDDIPF tip.

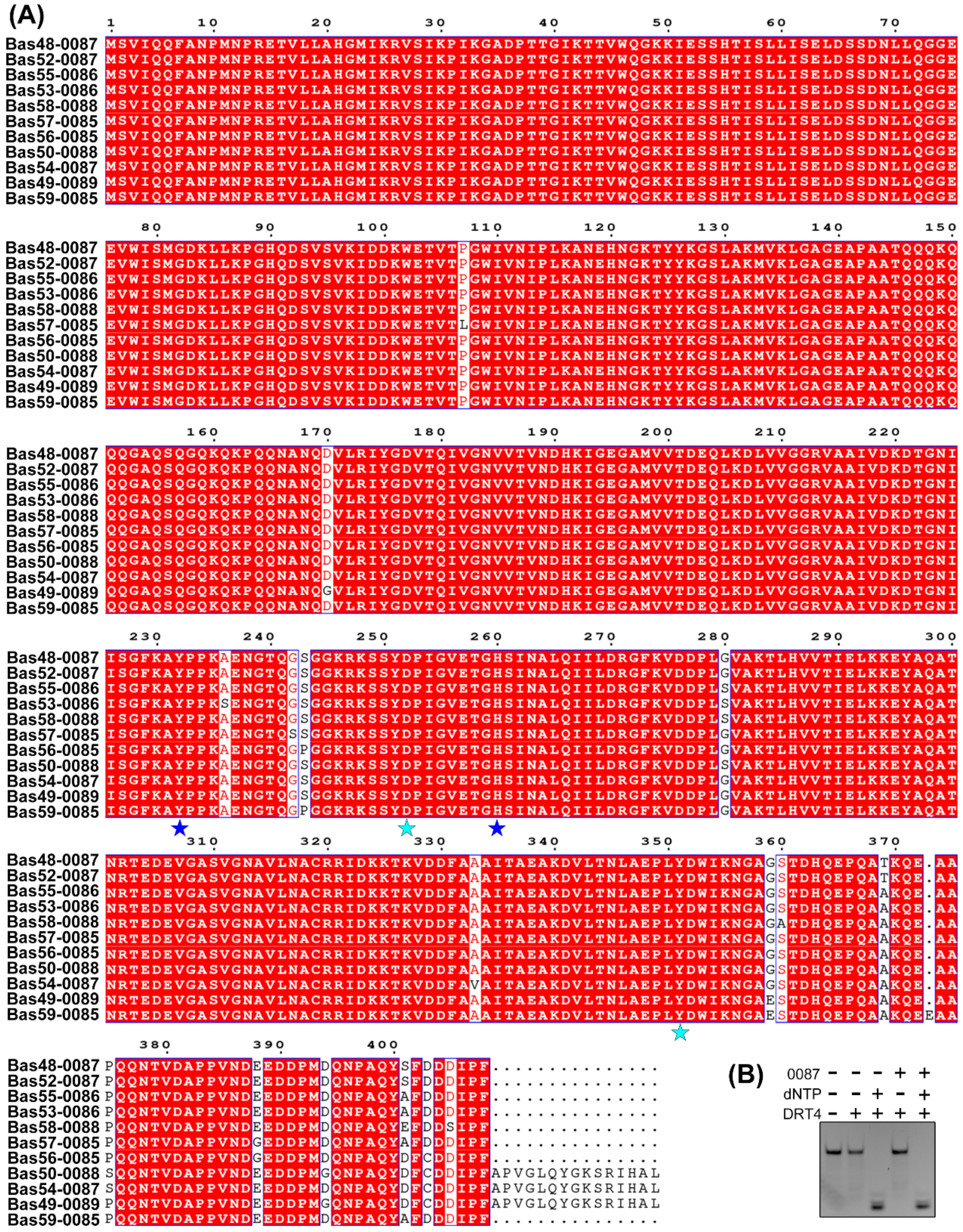


**Figure S5 | Bas48-0087 is conserved across a lot of related phages.**

**(A)** Multiple sequence alignment of QXV84166.1 (hypothetical protein bas52_0087), QXV75636.1 (hypothetical protein bas56_0085), QXV77554.1 (hypothetical protein bas50_0088), QXV77779.1 (hypothetical protein bas59_0085), QXV77991.1 (hypothetical protein bas49_0089), QXV79874.1 (hypothetical protein bas58_0088), QXV80986.1 (hypothetical protein bas55_0086), QXV81209.1 (hypothetical protein bas53_0086), QXV82607.1 (hypothetical protein bas54_0087), QXV82826.1 (hypothetical protein bas57_0085). The conserved residues are indicated with red shading. The mutant sites in Bas48-0087 were labeled with blue star and the mutant sites in Bas52-0087 were labeled with cyan star.

**(B)** ssRNA cleavage assays of DRT4 in the absence and presence of dNTPs and 0087.

**Table S1: Cryo-EM data collection, refinement and validation statistics**

| | DRT4-ssDNA |
|---|---|
| **Data collection and processing** | |
| Magnification | 96,000 |
| Voltage (keV) | 300 |
| Electron exposure ($e^{-}$/Å$^2$) | 50 |
| Defocus range (μm) | 1.5 to 2.5 |
| Pixel size (Å) | 0.838 |
| Symmetry imposed | C1 |
| Initial particle images (no.) | 824,252 |
| Final particle images (no.) | 385,016 |
| Map resolution (Å) | 2.65 |
| FSC threshold | 0.143 |
| Map resolution range (Å) | 1.6-999 |
| **Refinement** | |
| Initial model used (PDB code) | *ab-initio* |
| Model resolution (Å) | 1.51 |
| FSC threshold | 0.5 |
| Map sharpening *B* factor (Å$^2$) | -105.5 |
| Map Correlation Coefficient | 0.90 |
| **Model composition** | |
| Non-hydrogen atoms | 26,136 |
| Protein residues | 3,108 |
| Nucleotides | 30 |
| ***B* factor (Å$^2$)** | |
| Protein | 36.57 |
| Nucleotides | 32.39 |
| **R.m.s. deviations** | |
| Bond lengths (Å) | 0.008 |
| Bond angles (°) | 1.305 |
| **Validation** | |
| MolProbity score | 1.41 |
| Clash score | 3.30 |
| Poor rotamers (%) | 2.39 |
| **Ramachandran plot** | |
| Favored (%) | 98.63 |
| Allowed (%) | 1.37 |
| Disallowed (%) | 0.00 |
| **EMDB** | EMD-82404 |
| **PDB** | 43YZ |
